# Rescuing behavioral flexibility in a mouse model for OCD by enhancing reward-cue salience

**DOI:** 10.1101/2024.07.25.604103

**Authors:** Bastijn J.G. van den Boom, Sara De Carlo, Jennifer van Klaveren, Damiaan Denys, Ingo Willuhn

## Abstract

Deficits in cognitive flexibility are a frequent symptom of obsessive-compulsive disorder (OCD) and have been hypothesized to underlie compulsive behavior. Sign- and goal-tracking behaviors are thought to be related to cognitive flexibility, yet have not been studied in this context. To investigate the relationship between sign- and goal-tracking behavior and cognitive flexibility, we tested SAPAP3 knockout mice (SAPAP3^-/-^) and wild-type littermate controls in a Pavlovian reversal-learning task with two conditioned stimuli, one predicting reward delivery and the other reward omission. SAPAP3^-/-^ displayed a heterogenous reversal-learning performance: Half of the population failed to acquire the reversed cue-reward contingencies, whereas the other half reversed their approach behavior similar to control mice. Surprisingly, such behavioral inflexibility and compulsive-like grooming were unrelated, suggesting a non-causal relationship between these traits. Importantly, compromised reversal learning in impaired mice was associated with diminished sign-tracking behavior (and therefore presumably with an overreliance on goal-tracking behavior). Administration of the selective serotonin reuptake inhibitor (SSRI) fluoxetine, the first-line pharmacological OCD treatment, ameliorated both anxiety-like behavior and compulsive-like grooming, but did not improve behavioral flexibility in SAPAP3^-/-^. In contrast, enhancing reward-cue salience by altering conditioned stimuli brightness improved behavioral flexibility through augmenting sign-tracking behavior. These findings suggest that deficits in behavioral flexibility are associated with imbalanced sign- and goal-tracking behaviors in SAPAP3^-/-^, and enhancing reward-cue salience can rescue behavioral flexibility by restoring the balance. Thus, sign- and goal-tracking behavior might be an underexplored cognitive mechanism that could potentially be exploited to improve cognitive flexibility in OCD patients.

## Introduction

Obsessive-compulsive disorder (OCD) is a psychiatric disorder that affects 1-3% of the population and is characterized by intrusive thoughts (obsessions) and repetitive behaviors (compulsions), often associated with cognitive deficits^1–5^. Deficits in cognitive flexibility are thought to be a potential mechanism underlying the inability of patients to cease compulsive behaviors^6–9^, and if so, may be exploited as a potential target for therapeutic intervention. Previous studies examining cognitive flexibility in OCD have produced inconsistent results, with some studies demonstrating deficits^10–14^, whereas others did not^15–17^. These inconsistencies could be explained by differences in behavioral tasks used to probe cognitive flexibility and the large interindividual variability in cognitive flexibility of OCD patients^18^. Notably, a large meta-analysis confirmed the connection between OCD and impaired cognitive flexibility^8^.

The most basic form of behavioral flexibility can be investigated using a procedure called autoshaping^19,20^. During autoshaping, a conditioned stimulus (CS) (e.g., a cue light) triggers approach behavior, either to the CS itself (sign tracking) or to the location (goal tracking) where the unconditional stimulus is delivered (e.g., food reward)^21,22^, thereby probing different cognitive mechanisms underlying the behavior. Goal-tracking behavior, similar to model-based learning, is thought to rely on a cognitive map of the task to guide actions and is dependent on action-outcome associations^23,24^. In contrast, sign-tracking behavior is thought to be rooted in the CS acquiring incentive salience^25^, a process consistent with model-free learning^24^. Importantly, both strategies are in competition, and one strategy might dominate depending on the task requirements. After acquisition of the task, the cue-reward contingencies can be reversed to assess Pavlovian reversal learning. The previously rewarded CS (CS+) becomes the non-rewarded CS (CS-), and vice versa. Both model-free and model-based strategies, which are comparable to sign and goal tracking, respectively, are thought to be involved in successful reversal learning^26–28^. Our recent work provided evidence for the role of sign and goal tracking in reversal learning, where a lack of sign tracking coincided with poor reversal performance^29^.

The activity of serotonin neurons has been associated with behavioral flexibility, both in animals and humans^30,31^: Blocking serotonin reuptake^32^, depletion of serotonin^33–35^, and inhibiting serotonin neurons^36^ resulted in altered behavioral flexibility. Importantly, acute depletion of serotonin or tryptophan (a serotonin precursor) induced deficits in behavioral flexibility^33,34,37^, which could be reversed by selective serotonin reuptake inhibitor (SSRI) treatment in animals^35,38^. SSRI treatment enhanced cognitive flexibility in some human studies^39^, whereas others found impaired cognitive flexibility^40^. Interestingly, selective serotonin depletion^41^ and altered serotonin function in inbred mice^42^ resulted in enhanced sign-tracking behavior, although others found no effect^43^.

Preclinical animal models are a valuable tool to assess OCD-related symptoms, and the SAPAP3 mutant mouse model (SAPAP3^-/-^) is, arguably, the best-established genetic mouse model for OCD, exhibiting compulsive-like grooming, anxiety-like behavior^29,44^, altered habit formation^45,46^, and cognitive deficits^18,29,47^. In addition, these mice respond well to pharmacotherapy^44,48^ and deep-brain stimulation^49,50^, and, similar to OCD patients, symptoms are dependent on cortico-striatal functioning^51^.

Here, we used SAPAP3^-/-^ and their wild-type littermates (WT) to assess behavioral flexibility and examine the role of sign- and goal-tracking behavior that may underlie reversal learning. We altered animals’ preference to employ sign- or goal-tracking behavior by changing reward-cue salience^52^. Increasing the brightness of the CS or reducing background light (i.e., dimming the house light) enhances reward-cue salience and induces more sign-tracking behavior, which is dependent on bottom-up, feedforward input^53^. Finally, we assessed the effects of SSRI treatment on behavioral flexibility.

## Methods and Materials

### Subjects

Three cohorts of male and female SAPAP3^-/-^ (n=23, average age five months (range 3.5 - 6 months)) and their wild-type littermates (WT, n=25, average age five months (range 3.5 - 6.5 months) were trained on a Pavlovian reversal-learning task. A subset of these mice were tested in the open field for grooming behavior (SAPAP3^-/-^, n=15; WT, n=17) and the elevated plus maze (EPM) to examine anxiety (SAPAP3^-/-^, n=14; WT, n=15) during SSRI treatment. An additional group of mice (SAPAP3^-/-^, n=17; WT, n=14) were tested on the EPM without SSRI treatment. Animals were housed solitarily and food restricted to 85% of their free-feeding body weight. Mice were kept under a 12-hour reversed light/dark cycle with ad-libitum access to water and all behavioral procedures were performed during the dark phase. All experiments were in accordance with Dutch and European laws and approved by the Animal Experimentation Committee of the Royal Netherlands Academy of Arts and Sciences.

### Open-field grooming

Grooming was assessed in an open field (30x30x40 cm) located in a light-shielded chamber in the dark for 60 min. Overhead videos were recorded using Bonsai^54^, and analyzed using MouseTracker^55^ and a trained Janelia Automatic Animal Behavior Annotator^56^ (for methodological details of the classifier, see^57^).

### Elevated plus maze

Anxiety was assessed in the EPM, which consisted of two “open” and two “closed” arms (length 31 cm, width 8 cm) of non-transparent matte-white Trespa material located in a light-shielded chamber in the dark. Animals were video recorded for 10 minutes and data analyzed using Bonsai (for methodological details of EPM recordings, see^29^).

### Pavlovian reversal learning

#### Apparatus

Behavioral training and testing were performed in operant chambers (Med Associates Inc., 21.6 x 17.8 x 12.7 cm) equipped with cue lights flanking a reward magazine and a house light (40 W) (Figure 1A). Cue lights were custom-made to emit regular (2 mA) or enhanced (50 mA) light. Operant chambers were equipped with an infra-red beam on the opposite side of the reward magazine for the animals to initiate trials. Custom-written scripts in MED IV (Med Associates Inc.) were used to control task events. In addition, top-down videos were recorded (Sony CCD, 0.5 MP) and analyzed using Bonsai to examine animals’ responses during cue presentations.

**Figure 1.**
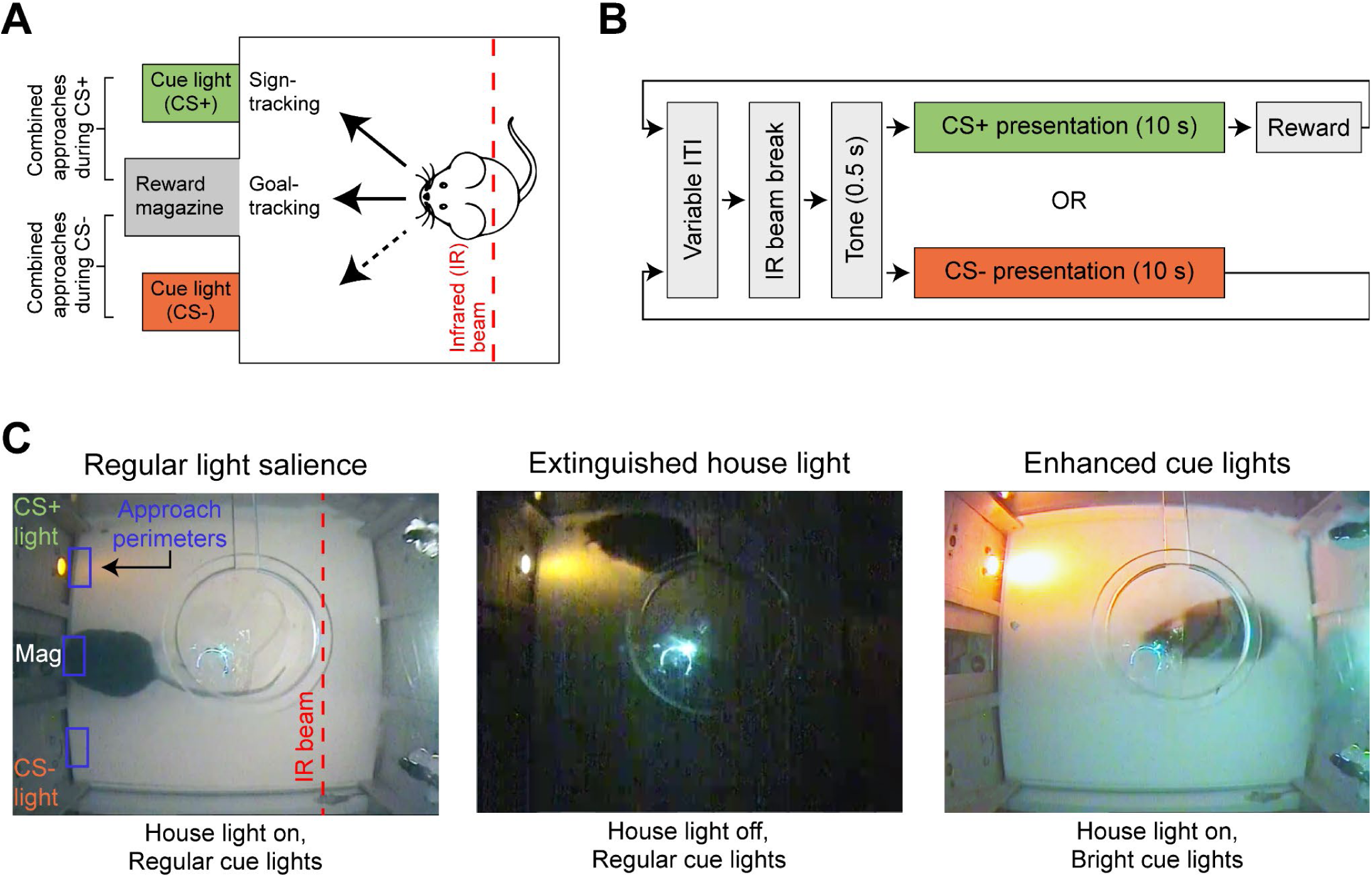
Experimental approach to examine behavioral flexibility across different cue salience conditions. **(A)** Schematic of top-down view of the operant chamber. A mouse could break an infrared beam located opposite to the cues and reward magazine to initiate a trial, which resulted in a CS+ or CS-presentation. Approaches towards the cues and magazine during CS+ and CS-presentations were recorded to assess whether animals differentiated between the CS+ and CS-. Time spent at the cue and magazine perimeter during CS+ presentations was used as a read-out for sign and goal tracking, respectively. **(B)** Schematic of the task design. After a variable inter-trial interval (ITI), animals were required to interrupt an infrared beam, which produced a short (0.5 s) auditory tone, followed by a 10 s CS+ or CS-presentation. CS+ was immediately followed by a reward (sucrose pellet), while CS- was never followed by a reward **(C)** Different light-salience conditions were used across three different experiments: Regular light-salience experiment (house light illuminated and regular cue lights), extinguished house-light experiment (house light extinguished and regular cue lights), and enhanced cue-light experiment (house light illuminated and intensity of cue lights enhanced). Video camera recordings were used to analyze the animals’ behavior.

#### Training

Training was based on an autoshaping protocol published previously^20^. Briefly, after being put on food restriction, animals received 20 rewards (20 mg sucrose pellets, Bio-Serv) in their homecage on two consecutive days. Next, the animals received two days of magazine training, during which they received 20 rewards on a fixed interval schedule of 60 s. Pavlovian training consisted of 40 trials per session, with 20 trials being CS+ and 20 trials being CS- (in randomized order within each session), and was conducted five days per week. After a variable inter-trial interval of 25 s (range: 10-40 s), mice had to interrupt the infrared beam at the back of the camber to initiate a trial. Trial initiation was signaled by an auditory tone (0.5 s click), followed by immediate visual presentation of the left or right cue light (CS+ and CS- location counterbalanced across animals) for 10 s. Independent of animals’ behavior, a CS+ trial produced a reward delivery, while a CS-trial would not be rewarded (Figure 1B). Besides position, the CSs were indistinguishable. After 17 sessions, cue-reward contingencies were reversed (previous CS+ cue light became CS- and vice versa). During the regular light-salience experiment, a re-reversal occurred after another 17 sessions. After reversal training (17 sessions), mice underwent a different reward-cue light salience experiment. The sequence of light salience experiments was counterbalanced between cohorts of mice (Figure 1C). Eventually, all cohorts received SSRI treatment during another regular light-salience experiment.

#### Exclusion and learning criterion

For each experiment, we excluded animals that did not associate the CS+ with reward (based on the last five sessions before reversal). This learning criterion included: 1) more CS+ approaches than CS- and 2) CS+ approaches (cue and magazine) in at least 70% of the trials. This allowed us to include animals that showed behavioral preference for the CS+ compared to the CS- and were engaged in the task. We employed the same learning criterion to define animals as “impaired” who did not fulfill this learning criterion after reversal (based on the last five sessions after reversal). This criterion was predefined and used in our previous work^29^.

#### Performance measures

Animals’ behavior during CS presentations was analyzed based on video recordings by measuring animals’ presence in the task-relevant approach perimeters (Figure 1A). In particular, we measured 1) number of trials with an approach towards cue and magazine (binary measure), 2) total number of approaches toward cue and magazine (total measure), and 3) time spent at cue or magazine during CS+ (time measure). The binary measure was used for exclusions and assessment of learning criteria; the total measure was used to determine if the animals differentiated between the CS+ and CS-, and the time measure was used to distinguish sign- and goal-tracking behavior.

#### SSRI application

To examine whether the SSRI treatment improved behavioral flexibility, animals received daily intraperitoneal injections of 5 mg/kg of fluoxetine hydrochloride (fluoxetine, LKT Laboratories Inc.) after being trained on the Pavlovian task. Animals received five days of SSRI injections, then five days of Pavlovian training combined with SSRI treatment, followed by reversal training for another 17 days combined with SSRI application. After reversal training, animals were tested on the open field and EPM for SSRI-induced altered grooming and anxiety-like behavior, respectively.

#### Light salience experiments

We assessed behavioral flexibility under different reward-cue salience conditions across three different experiments: Regular light-salience experiment (house light illuminated and regular cue lights), extinguished house-light experiment (house light extinguished and regular cue lights), and enhanced cue-light experiment (house light illuminated and light intensity of both cue lights enhanced). The regular light-salience experiment made use of the standard Med Associates house light and cue lights.

### Statistical analyses

We employed paired and independent *t*-tests, one- or two-way ANOVAs followed by post-hoc multiple comparison tests, Pearson correlations, and chi-square tests. The *t*-tests were corrected for multiple comparisons using the Holm-Bonferroni method^58^ and the post-hoc tests using Tukey-Kramer tests. Statistical significance was defined as *p* < 0.05. All statistical analyses were performed using Matlab (R2016b and R2020b, MathWorks Inc.). Data are presented as mean + SEM.

## Results

### SAPAP3^-/-^ show impaired behavioral flexibility associated with imbalanced sign- and goal-tracking behavior

As previously reported, SAPAP3^-/-^ initiated more grooming bouts and spent more time grooming compared to WT (Figure S1A). We trained SAPAP3^-/-^ and WT on a Pavlovian reversal-learning task to replicate previously reported behavioral inflexibility in SAPAP3^-/-18,29,47^ and examine its relation to sign- and goal-tracking behavior. To infer overall task performance independent of sign- or goal-tracking strategies, approaches towards the cue and magazine during CS+ or CS-presentation were combined into a single measure of combined approaches. During the regular light-salience experiment, we found no differences in the learning rate nor session duration between WT and SAPAP3^-/-^ (Figure S1B,C), suggesting similar learning capabilities and motivational drive between genotypes, respectively. WT displayed differentiation between CS+ and CS- (CS: F_1,14_=136.40, *p*<0.001; reversal: F_2,28_=0.67, *p*=0.519; interaction: F_2,28_=2.01, *p*=0.153) before reversal (*p*<0.001), after reversal (*p*<0.001), and after re-reversal (*p*<0.001) (Figure 2A). Collectively, the SAPAP3^-/-^ population differentiated between CS+ and CS- (CS: F_1,11_=32.16, *p*<0.001; reversal: F_2,22_=0.09, *p*=0.911; interaction: F_2,22_=0.81, *p*=0.458) before reversal (*p*<0.001), after reversal (*p*=0.009), and after re-reversal (*p*=0.002) (Figure 2B). However, applying our learning criteria showed that 50% of the SAPAP3^-/-^ failed to acquire the new cue-reward contingencies, compared to only 6% of the WT (*X*^2^_(1, N=28)_=7, *p*=0.008) (Figure 2C). Indeed, when splitting the SAPAP3^-/-^ population into animals that successfully acquired the new cue-reward contingencies (reversing SAPAP3^-/-^) and those that failed to do so (impaired SAPAP3^-/-^), we found that reversing SAPAP3^-/-^ were able to learn the task (CS: F_1,5_=118.60, *p*<0.001; reversal: F_1,5_=2.87, *p*=0.151; interaction: F_1,5_=1.68, *p*=0.252) before (*p*=0.008) and after (*p*<0.001) reversal (Figure 2D), while impaired SAPAP3^-/-^ only acquired the task (CS: F_1,5_=23.05, *p*=0.005; reversal: F_1,5_=2.60, *p*=0.168; interaction: F_1,5_=9.44, *p*=0.028) before reversal (*p*=0.006) but failed to update their approach behavior after reversal (*p*=0.937) (Figure 2E). Importantly, reversing SAPAP3^-/-^ and impaired SAPAP3^-/-^ did not differ in the amount of grooming, learning rate of the task, or session duration (Figure S1D,E,F), suggesting that impaired SAPAP3^-/-^ specifically lack behavioral flexibility. In addition, the amount of grooming in the open field did not correlate with task performance before or after reversal (Figure S1G,H).

**Figure 2.**
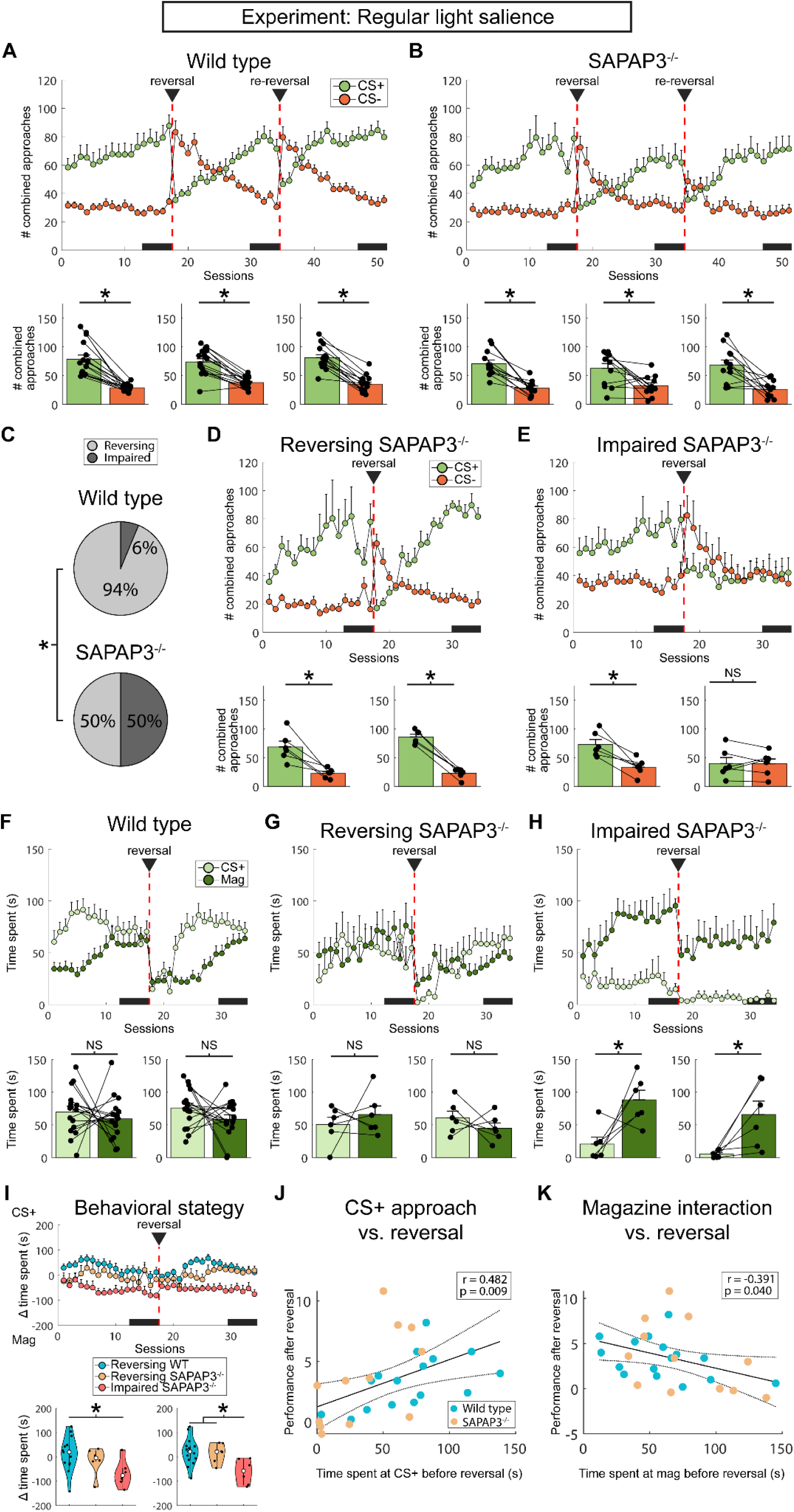
Impaired SAPAP3^-/-^ failed to adjust behavior after changed cue-reward contingencies, accompanied by imbalance sign- and goal-tracking behavior. **(A)** Top: Total number of approaches toward cue and magazine during CS+ (green) and CS- (red) trials for WT (n=16). Bottom: Quantification of differentiation between CS+ and CS- before reversal (left), before re-reversal (middle), and after re-reversal (right). Data points are individual animals. WT successfully adjusted their behavior after reversal and re-reversal. **(B)** Similar to (A), except for SAPAP3^-/-^ (n=12). Across the entire SAPAP3^-/-^ population, mice were able to reverse and re-reverse their behavior upon altering cue-reward contingencies (top). Quantification of the reversal (bottom, middle panel) revealed two populations: One population showed differentiation between CS+ and CS-(reversing), and the other did not (impaired). **(C)** We quantified the percentage of reversing and impaired mice and found that 50% of the SAPAP3^-/-^ showed behavioral deficits compared to 6% of the WT. **(D)** SAPAP3^-/-^ that met reversal criteria differentiated between CS+ and CS- before and after reversal (“reversing SAPAP3^-/-^”, n=6) based on total number of approaches (top), as quantified below (bottom). **(E)** “Impaired SAPAP3^-/-^“ (n=6) differentiated between CS+ and CS- before reversal but not after (top), as indicated by bar graphs (bottom). **(F)** WT showed balanced sign- and goal-tracking behavior before and after reversal, as indicated by time spent at the CS+ (light green) and magazine (dark green). Quantification (bottom) revealed no difference between CS+ and magazine time spent. **(G)** Similar to WT, the reversing SAPAP3^-/-^ group demonstrated balanced sign- and goal-tracking behavior (top), both before and after reversal (bottom). **(H)** Impaired SAPAP3^-/-^ primarily employed a goal-tracking strategy (top), which was present before reversal and and after reversal (bottom)l. **(I)** We calculated a difference score by subtracting time spent at the magazine from CS+. Impaired SAPAP3^-/-^ consistently show enhanced goal tracking (top). Quantifying the difference in behavioral strategy between WT, reversing SAPAP3^-/-^, and impaired SAPAP3^-/-^ revealed enhanced goal tracking of impaired SAPAP3^-/-^ before and after reversal (bottom). **(J)** Correlation analyses revealed a positive relationship between sign-tracking before reversal and performance after reversal across all mice (n=28). **(K)** Goal-tracking before reversal was negatively correlated with performance after reversal (n=28). \**p* < 0.05, NS = not significant.

We used the time spent in the approach perimeter (Figure 1C) of the cue and magazine during CS+ presentation as a measure for sign and goal tracking, respectively. WT displayed balanced sign- and goal-tracking strategies before and after reversal (strategy: F_1,15_=0.98, *p*=0.337; reversal: F_1,15_=1.05, *p*=0.323; interaction: F_1,15_=0.34, *p*=0.571) (Figure 2F). Similar to WT, reversing SAPAP3^-/-^ employed a balanced sign- and goal-tracking strategy before and after reversal (strategy: F_1,5_=0.01, *p*=0.978; reversal: F_1,5_=0.58, *p*=0.480; interaction: F_1,5_=0.91, *p*=0.384) (Figure 2G). In contrast to WT and reversing SAPAP3^-/-^, impaired SAPAP3^-/-^ strongly relied on goal-tracking behavior (strategy: F_1,5_=33.32, *p*=0.002; reversal: F_1,5_=2.43, *p*=0.180; interaction: F_1,5_=0.03, *p*=0.863), both before (*p*=0.032) and after (*p*=0.036) reversal (Figure 2H). Directly comparing the three groups revealed significant differences before (genotype: F_2,24_=3.69; *p*=0.040; post hoc WT vs impaired SAPAP3^-/-^ *p*=0.032) and after (genotype: F_2,24_=5.70; *p*=0.009; post hoc WT vs impaired SAPAP3^-/-^: *p*=0.008; reversing SAPAP3^-/-^ vs impaired SAPAP3^-/-^: *p*=0.044) reversal (Figure 2I). Finally, as shown previously^29^, sign tracking after reversal positively correlated with reversal performance (Figure S1I), although goal tracking did not correlate with reversal performance (Figure S1J). Importantly, we found a positive correlation between sign tracking before reversal and performance after reversal (r=0.48, *p*=0.009) (Figure 2J) and a negative correlation between goal tracking before reversal and performance after reversal (r=-0.39, *p*=0.040) (Figure 2K), suggesting that the employed behavioral strategy before the reversal occurs predicts the degree of behavioral flexibility. Taken together, half of the SAPAP3^-/-^ population displayed behavioral inflexibility, accompanied by over-reliance on goal-tracking behavior.

### Chronic SSRI treatment does not alter behavior inflexibility

To investigate the effects of chronic SSRI treatment on behavioral flexibility, animals received daily fluoxetine injections for one month. Comparing compulsive-like grooming before and after chronic SSRI application revealed an effect of genotype and treatment (genotype: F_1,59_=22.44, *p*<0.001; treatment: F_1,59_=6.96, *p*=0.011; interaction: F_1,59_=1.19, *p*=0.280) (Figure 3A). While WT did not show a significant reduction in grooming (*p*=0.051), SAPAP3^-/-^ did (*p*=0.026). By comparing SSRI-treated animals against not-treated animals tested on the EPM (genotype: F_1,56_=8.69, *p*=0.005; treatment: F_1,56_=33.27, *p*<0.001; interaction: F_1,56_=0.67, *p*=0.417) (Figure 3B), we found reduced anxiety-like behavior in both WT (*p*=0.005) and SAPAP3^-/-^ (*p*<0.001). Next, we examined behavioral flexibility during chronic SSRI treatment. WT were able to adapt their behavior successfully (CS: F_1,12_=43.80, *p*<0.001; reversal: F_1,12_=3.61, *p*=0.082; interaction: F_1,12_=24.56, *p*<0.001), indicated by CS+ and CS-discrimination before (*p*<0.001) and after (*p*=0.001) reversal (Figure 3C). In contrast, SAPAP3^-/-^ were not able to adapt their behavior (CS: F_1,11_=26.93, *p*<0.001; reversal: F_1,11_=4.98, *p*=0.048; interaction: F_1,11_=8.20, *p*=0.015), illustrated by CS+ and CS-discrimination before (*p*<0.001), but not after (*p*=0.059) reversal (Figure 3D). Directly comparing the fraction of animals that learned the reversal revealed significant differences between WT and SAPAP3^-/-^ (*X*^2^_(1, *N*=25)_=5.00, *p*=0.025) (Figure 3E). Dividing the SAPAP3^-/-^ population into successful reversing and impaired groups revealed that reversing SAPAP3^-/-^ (CS: F_1,4_=23.93, *p*=0.008; reversal: F_1,4_=0.90, *p*=0.396; interaction: F_1,4_=2.11, *p*=0.220) learned the task before (*p*=0.026) and after (*p*<0.001) reversal (Figure 3F), while impaired SAPAP3^-/-^ (CS: F_1,6_=10.29, *p*=0.018; reversal: F_1,6_=4.05, *p*=0.091; interaction: F_1,6_=6.13, *p*=0.048) differentiated between CS+ and CS- before reversal (*p*=0.014), but not after (*p*=0.826) (Figure 3G). Together, these data demonstrate that SSRI treatment reduces compulsive- and anxiety-like behaviors, but is ineffective in rescuing behavioral flexibility.

**Figure 3.**
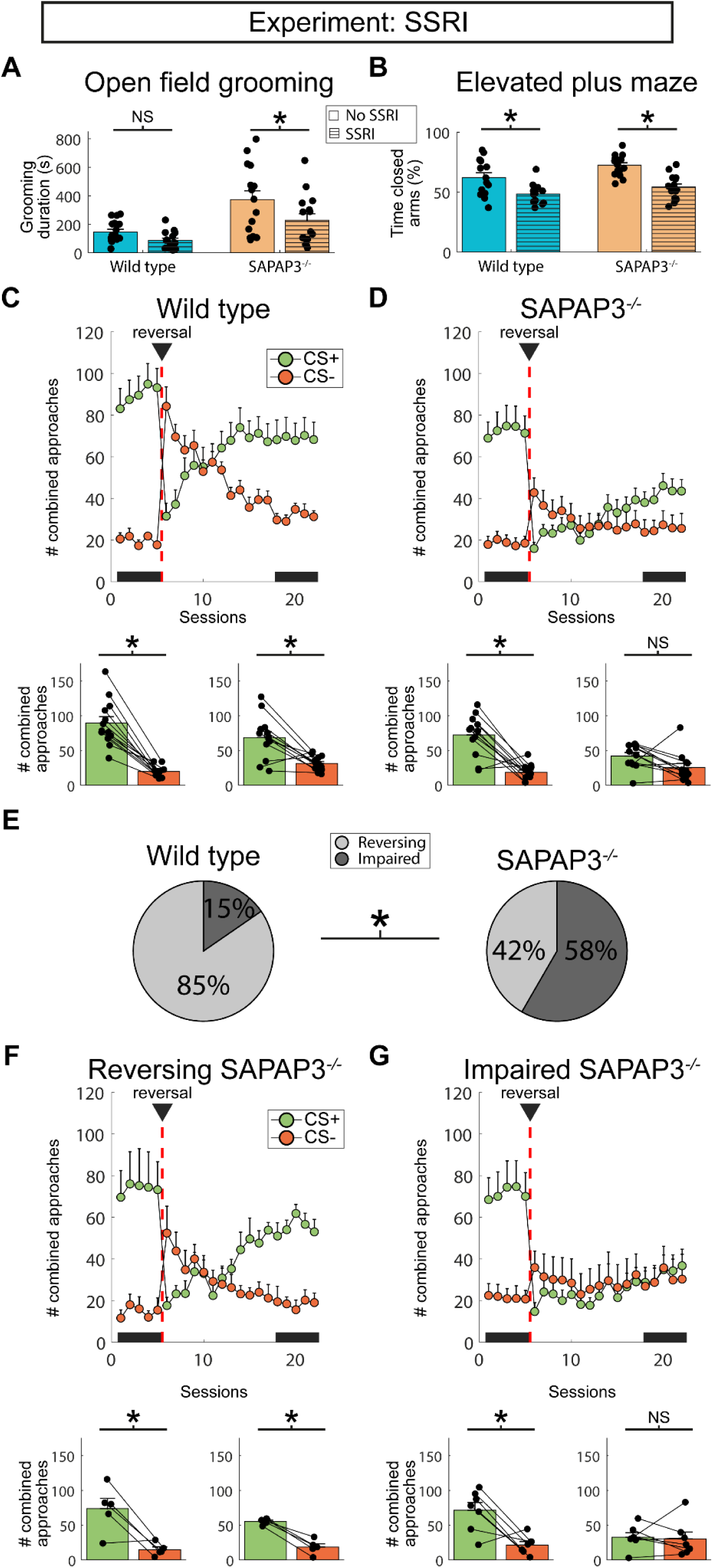
SSRI treatment did not rescue behavioral flexibility. **(A)** SSRI treatment significantly reduced compulsive-like grooming in SAPAP3^-/-^ (n=15), without affecting grooming in WT (n=17). Data points are individual animals, repeated before and after SSRI treatment. **(B)** Both WT (no SSRI n=14, SSRI n=15) and SAPAP3^-/-^ (no SSRI n=17, SSRI n=14) exhibited reduced anxiety-like behavior following SSRI treatment. Each data point is an individual animal. **(C)** The number of combined approaches (to the cues and magazine) revealed that WT (n=13) injected with SSRI were able to reverse their behavior (top), as quantified before and after reversal (below). **(D)** SSRI treatment did not prevent SAPAP3^-/-^ (n=12) from differentiating between CS+ and CS- before reversal, but selectively impaired reversal learning (top), as quantified below. **(E)** Quantifying the percentage of reversing and impaired animals revealed that the majority of SAPAP3^-/-^ (58%) was unable to reverse their behavior, which was significantly different from WT (15%). Splitting the data into reversing (G) and impaired (F) SAPAP3^-/-^ showed that reversing SAPAP3^-/-^ (n=5) were able to alter their behavior upon changing cue-reward contingencies, while impaired SAPAP3^-/-^ (n=7) were unable to do so. \**p* < 0.05, NS = not significant.

### Enhanced reward-cue salience rescues behavioral flexibility by altering behavioral strategy

Impaired SAPAP3^-/-^ overly rely on a goal-tracking strategy, a behavior that negatively correlated with reversal performance. Previous work has suggested that increasing reward-cue salience improves sign-tracking behavior^52^. Therefore, in two separate experiments, we either extinguished the house light or enhanced the cue lights to improve reward-cue salience. We hypothesized that increasing reward-cue salience could promote a more balanced engagement of sign- and goal-tracking behaviors, potentially leading to improved reversal learning in SAPAP3^-/-^. During the extinguished house-light experiment, WT were able to discriminate CS+ and CS- (CS: F_1,14_=196.43, *p*<0.001; reversal: F_1,14_=0.79, *p*=0.390; interaction: F_1,14_=4.74, *p*=0.047), both before (*p*<0.001) and after (*p*<0.001) reversal (Figure 4A). Similarly to WT, SAPAP3^-/-^ discriminated between CS+ and CS- (CS: F_1,13_=29.44, *p*<0.001; reversal: F_1,13_=0.01, *p*=0.96; interaction: F_1,13_=0.99, *p*=0.337) before (*p*<0.001) and after (*p*=0.001) reversal (Figure 4B). Importantly, directly comparing the fraction of animals that learned the reversal revealed no differences between WT and SAPAP3^-/-^ (*X*^2^ *_N_*_=29)_=1.11, *p*=0.291) (Figure 4C). Similarly, we found no difference in the behavioral strategy employed by WT and SAPAP3^-/-^ before (t(26)=1.64, *p*=0.113) and after (t(26)=1.91, *p*=0.068) reversal (Figure 4D). Sign tracking before reversal did not correlate significantly with performance after reversal (r=0.33, *p*=0.082) (Figure 4E). Similarly to the regular light-salience experiment, goal tracking before reversal negatively correlated with performance after reversal (r=-0.55, *p*=0.002) (Figure 4F).

**Figure 4.**
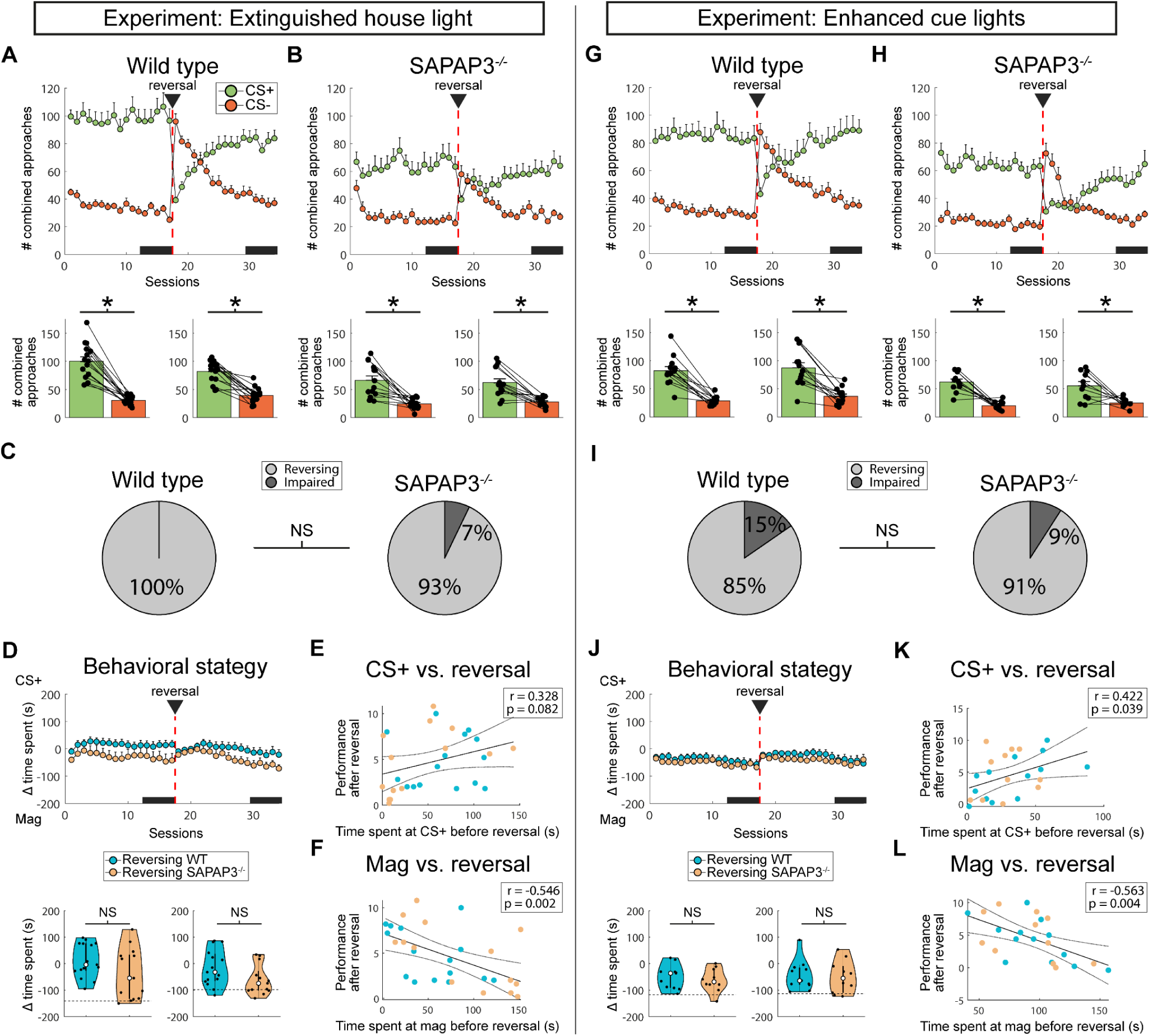
Altering cue salience restored behavior flexibility by balancing sign- and goal-tracking behavior. **(A)** WT (n=15) and **(B)** SAPAP3^-/-^ (n=14) were able to successfully reverse their approach when the house light was turned off (top). Quantification revealed that both genotypes were able to learn the task and change their behavior upon reversal (bottom). Data points represent individual animals. **(C)** The number of animals that acquired the reversal did not differ between WT (100%) and SAPAP3^-/-^ (93 %). **(D)** Both WT (blue) and SAPAP3^-/-^ (orange) employed a balanced sign- and goal-tracking strategy (top). The behavioral strategy employed by WT and SAPAP3^-/-^ did not differ significantly (bottom). Horizontal dotted lines (left and right) indicate the single SAPAP3^-/-^ that failed to acquire the reversal and relied on goal tracking. **(E)** Across all animals (n=29), sign-tracking before reversal did not significantly correlate with performance during reversal. **(F)** Goal-tracking before reversal was negatively correlated with reversal performance. Both **(G)** WT (n=13) and **(H)** SAPAP3^-/-^ (n=11) demonstrated reversal learning during the enhanced-cue experiment, similar to (A) and (B), respectively. **(I)** The number of animals that learned to adjust their behavior upon reversal did not differ between WT (85 %) and SAPAP3^-/-^ (91 %). **(J)** No difference in behavioral strategy employed by WT and SAPAP3^-/-^ was found (top). Similar to (D), time spent at the CS+ or magazine did not differ between WT and SAPAP3^-/-^ before and after reversal (bottom). The horizontal dotted line depicts the single SAPAP3^-/-^ that failed to update its behavior after reversal and relied on goal tracking. **(K)** Sign-tracking before reversal positively correlated with reversal performance across all animals (n=24). **(L)** Goal-tracking before reversal negatively correlated with performance during reversal. \**p* < 0.05, NS = not significant.

During the enhanced cue-light experiment, WT were able to differentiate between the cues (CS: F_1,12_=73.65, *p*<0.001; reversal: F_1,12_=1.37, *p*=0.264; interaction: F_1,12_=0.11, *p*=0.747) before (*p*<0.001) and after (*p*<0.001) reversal (Figure 4G). Again, SAPAP3^-/-^ also discriminated between the cues (CS: F_1,10_=30.44, *p*<0.001; reversal: F_1,10_=0.06, *p*=0.817; interaction: F_1,10_=2.20, *p*=0.169) before (*p*<0.001) and after (*p*=0.005) reversal (Figure 4H). Similar to the extinguished house-light experiment, we found no difference in the number of animals that learned the reversal between WT and SAPAP3^-/-^ (*X*^2^ =0.22, *p*=0.642) (Figure 4I). In addition, the behavioral strategy employed did not differ between WT and SAPAP3^-/-^ before (t(19)=0.61, *p*=0.546) or after (t(19)=0.24, *p*=0.811) reversal (Figure 4J). Sign tracking before reversal positively correlated with performance after reversal (r=0.42, *p*=0.040) (Figure 4K), while goal tracking before reversal negatively correlated with performance after reversal (r=-0.56, *p*=0.004) (Figure 4L). Collectively, these experiments demonstrate that increasing the reward-cue salience improves reversal learning in SAPAP3^-/-^ by altering the behavioral strategy employed.

### Impaired SAPAP3^-/-^ lack sign-tracking behavior

To directly compare behavioral strategies employed by the different groups of animals, we computed a difference score by subtracting time spent during CS- from CS+ in the approach perimeter (Figure 1C) at the cue and magazine. This difference score considers the preference for the CS+, which is indicative of how well animals differentiate between the cues. During the regular light-salience experiment, we found an effect of group on cue-approach behavior before (F_2,24_=4.40, *p*=0.024) and after reversal (F_2,24_=16.81, *p*<0.001) (Figure 5A), but not on magazine-approach behavior (before: F_2,24_=0.83, *p*=0.450; after: F_2,24_=3.29, *p*=0.055) (Figure 5B). Before reversal, impaired SAPAP3^-/-^ displayed a lack of sign tracking (WT vs impaired SAPAP3^-/-^: *p*=0.019), which persists after reversal (WT vs impaired SAPAP3^-/-^: *p*<0.001; reversing vs impaired SAPAP3^-/-^: *p*<0.001). During the SSRI experiment, we found similar results: An effect of group on cue-approach behavior before (F_2,20_=4.17, *p*=0.031) and after reversal (F_2,20_=9.42, *p*=0.001) (Figure 5C), and partially on magazine-approach behavior after reversal (F_2,20_=6.16, *p*=0.008) (Figure 5D). Similar to the regular light-salience experiment, impaired SAPAP3^-/-^ lacked sign-tracking behavior, both before (WT vs impaired SAPAP3^-/-^: *p*=0.025) and after reversal (WT vs impaired SAPAP3^-/-^: *p*=0.001; reversing vs impaired SAPAP3^-/-^: *p*=0.012s). After reversal, we found a suppression of goal-tracking behavior in both reversing (WT vs reversing SAPAP3^-/-^: *p*=0.041) and impaired SAPAP3^-/-^ (WT vs impaired SAPAP3^-/-^: *p*=0.015). Together, these data suggest that a lack of sign tracking before and after reversal is associated with deficits in reversal learning impaired SAPAP3^-/-^.

**Figure 5.**
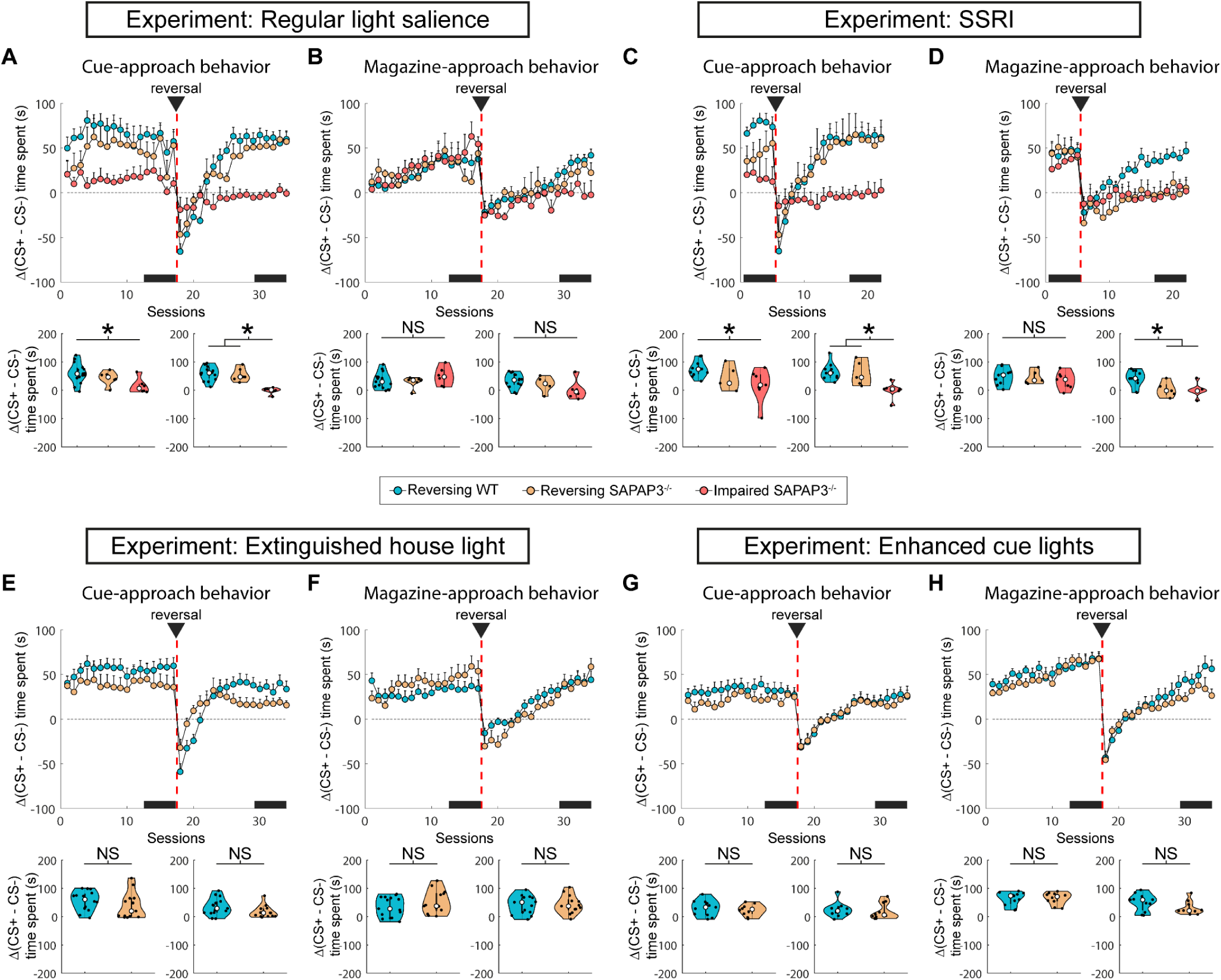
Impaired SAPAP3^-/-^ lack cue-approach behavior before and after reversal. Regular light-salience experiment: **(A)** Reduced cue-approach behavior (difference in time spent at cue during CS+ and CS-) in impaired SAPAP3^-/-^ (top), quantified before and after reversal (bottom). Each data point represents an individual animal. **(B)** Magazine-approach behavior (difference in time spent at the magazine during CS+ and CS-) was not affected. SSRI experiment: **(C)** Same as (A), impaired SAPAP3^-/-^ show reduced cue-approach behavior (top), both before and after reversal (bottom). **(D)** Same as B, although impaired SAPAP3^-/-^ showed significantly reduced magazine interaction after reversal (bottom). Extinguished house-light experiment: **(E)** No difference in cue-approach behavior was found between WT and SAPAP3^-/-^ (top). Quantification before and after reversal revealed no significant difference (bottom). **(F)** SAPAP3^-/-^ employed magazine-approach behavior to a similar degree as WT (top). No significant difference was found (bottom). Enhanced cue-light experiment: **(G)** same as (E), WT and SAPAP3^-/-^ employed cue-approach behavior to a similar degree (top) with no significant differences before or after reversal (bottom). **(H)** same as (F), magazine-approach behavior was comparable between WT and SAPAP3^-/-^ (top) with no significant differences (bottom). \**p* < 0.05, NS = not significant.

During the extinguished house-light experiment, we found no differences between WT and SAPAP3^-/-^ in cue-approach behavior (before: t(26)=1.21, *p*=0.237; after: t(26)=1.81, *p*=0.082) or magazine-approach behavior (before: t(26)=1.14, *p*=0.264; after: t(26)=0.11, *p*=0.915) (Figure 5E,F). Similarly, during the enhanced cue-light experiment, we found no differences between WT and SAPAP3^-/-^ in cue-approach behavior (before: t(19)=0.67, *p*=0.510; after: t(19)=0.44, *p*=0.668) or magazine-approach behavior (before: t(19)=0.15, *p*=0.879; after: t(19)=1.64, *p*=0.118) (Figure 5G,H). Taken together, these findings suggest that a lack of sign tracking might be involved in reversal-learning deficits, which can be altered by changing reward-cue salience.

## Discussion

We demonstrate that SAPAP3^-/-^, a widely used animal model for OCD, exhibit impairments in OCD-relevant behavioral flexibility using a Pavlovian reversal-learning task. We replicated our previous findings^29^ and provided ways to rescue these deficits. Approximately 50% of the SAPAP3^-/-^ failed to adjust their behavior upon reversal of cue-reward contingencies, whereas the other half performed indistinguishable from WT controls. Impaired SAPAP3^-/-^ and reversing SAPAP3^-/-^ differed in their display of balanced sign-tracking and goal-tracking behaviors, which was predictive of reversal-learning performance. To explore the relationship between balanced sign and goal tracking and reversal-learning performance, we altered reward-cue salience in two different ways, which resulted in employment of balanced behavioral strategies and reinstated reversal learning in impaired mice.

### Over-reliance on goal-tracking behavior could be related to goal-directed deficits in OCD

Compulsive behaviors in OCD patients are learned behaviors, which, over time and with repetition, eventually become stereotypical. Habits and compulsive behaviors are both performed independently of the outcome of their actions. Due to the similarities with habits, compulsivity has been described as exaggerated habitual responding^59^. Importantly, this could be caused by deficits in habit formation or expression, as well as deficits in goal-directed functioning. Recent work has suggested that OCD patients experience deficits in goal-directed control, resulting in overreliance on habits^60–62^. We report that impaired SAPAP3^-/-^ primarily display goal tracking, which is thought to be related to goal-directed behavior, since both behaviors depend on the outcome^24,63^. We speculate that impaired SAPAP3^-/-^ are “stuck” in a goal-tracking strategy and not able to cease this behavior, which could be interpreted as a goal-directed behavior deficit. Interestingly, previous work has shown that SAPAP3^-/-^ display altered reward processing and impaired habit formation^45,46^, which could be caused by overreliance on goal-directed behavior. Indeed, similar to our findings, Ehmer and colleagues found SAPAP3^-/-^ to over-rely on goal-directed behavior, even when habitual responding was promoted^45^ (although Hadjas and colleagues found opposite results^46^, potentially due to differences in behavioral task and age range of the mice). SAPAP3^-/-^ were unable to form habits, a process tightly linked to sign-tracking behavior^24^. These findings are consistent with clinical observations of OCD patients being “stuck” in a loop of goal-directed actions rather than habitual actions^64^. This is a crucial distinction rooted in the finding that, during compulsive behaviors, OCD patients are aware of their senseless behavior and actively select goal-directed actions that together make up their compulsive behavior^64^. Additionally, decision-making in OCD patients is accompanied by disrupted expected value and impaired knowledge of the outcome, indicating a deficit in goal-directed control^61,62^. Taken together, SAPAP3^-/-^ might over-rely on goal-tracking behavior, suggesting a deficit in goal-directed behavior.

### Motivational and sensory salience underlying improved behavioral flexibility

Using reward-cue salience, we improved behavioral flexibility in SAPAP3^-/-^. Importantly, we used two different methods to enhance reward-cue salience with similar effects on behavioral flexibility: Either by increasing the reward-cue light or by diminishing the house light (background light). By increasing the reward-cue lights, we improved, among others, sensory salience, which is described as conspicuous stimulus features^65^. The enhanced cue lights emit more photons, which could steer attention to the cues and allow for easier detection by the animal. On the contrary, diminishing the background house light does not change the amount of photons emitted by the cue lights. However, one could still argue that the increased contrast between cue lights and background does result in enhanced sensory salience^66^. One argument against sensory salience underlying restored behavioral flexibility is the lack of learning rate differences between reversing and impaired SAPAP3^-/-^, suggesting that both groups detect the cue lights equally well. In addition, we found no evidence for improved performance before reversal of cue-reward contingencies during extinguished house light or enhanced cue lights conditions compared to the regular light-salience condition, arguing that the detection of cue lights does not explain the lack of behavioral flexibility in impaired SAPAP3^-/-^. Therefore, motivational salience, which is a cognitive process that drives behavior towards or away from a particular stimulus or outcome, seems more likely to underlie enhanced reward-cue salience induced restored behavioral flexibility^67^. The attractive form of motivational salience, incentive salience, is responsible for approach behavior towards a desirable stimulus^25,67,68^. Importantly, for sign-tracking behavior, but not goal tracking, the reward-predictive cue is more attractive and acquires the properties of an incentive stimulus, resulting in approach behavior towards it^69–71^. Indeed, during the regular light-salience condition, when some of the properties of the incentive stimulus are potentially absent in the reward cue, impaired SAPAP3^-/-^ lack sign-tracking behavior. During reward-cue salience conditions, we found that SAPAP3^-/-^ employed both sign and goal tracking (similar to WT), suggesting that motivational salience has been restored. However, we cannot differentiate between motivational and sensory salience in our task without altering the reward outcome (e.g., increasing the amount of rewards per trial). Future work should use variable reward sizes or a probabilistic version of the task to probe the difference between motivational and sensory salience.

### Balanced sign tracking and goal tracking is needed for reversal learning

During the acquisition of the task, impaired SAPAP3^-/-^ employed a different behavioral strategy compared to WT and reversing SAPAP3^-/-^. In anticipation of reward (during CS+ presentation), both WT and reversing SAPAP3^-/-^ approached both the CS+ location (sign tracking) and the reward magazine (goal tracking), whereas impaired SAPAP3^-/-^ almost exclusively approached the reward magazine (goal tracking). We found that both sign and goal tracking before reversal predicted reversal-learning performance, suggesting that balanced sign- and goal-tracking strategies are needed for optimal reversal learning. Interestingly, in our previous work^29^, we found that SAPAP3^-/-^ relied entirely on goal tracking before reversal, which resulted in the majority of SAPAP3^-/-^ failing to update their behavior upon reversal of cue-reward contingencies. We hypothesize that the cues used in these works (relatively bright cue lights in the current experiments vs dim touch screen illumination in our previous work) might explain the differences in behavioral strategies employed by SAPAP^-/-^. Sign-tracking behavior is believed to rely on the CS+ acquiring incentive salience^25^, while goal-tracking behavior is unrelated to incentive motivation^23^. It has been hypothesized that successful reversal learning depends on knowledge of reward history from previous choices combined with a cognitive map of the task to guide animals’ future choices^26^, which could be cognitive mechanisms expressed by sign and goal tracking^30^, respectively. However, it is thought that these processes are necessary under conditions where cue-reward contingencies change with high frequency, resulting in animals tracking whether they received a reward or not and updating their belief state that reversals can occur^27,28^. Interestingly, these results have been reported for deterministic^27^ and probabilistic^28^ reversal-learning tasks, which vary in their degree of complexity. In our deterministic task, animals experience low cue-reward contingency reversals, which correlated with balanced sign and goal tracking. To probe the necessity of balanced strategies on behavioral flexibility, we tested whether enhanced reward-cue salience would induce sign tracking in the SAPAP3^-/-^ population (which primarily relied on goal tracking and lacked sign tracking under regular-light salience conditions) and would result in more reversing SAPAP3^-/-^ animals. Indeed, altering reward-cue salience resulted in more balanced sign- and goal-tracking behavior, as previously shown^52^. Notably, balanced sign- and goal-tracking strategy coincided with more animals displaying successful reversal learning. However, we cannot exclude other (indirect) mechanisms underlying the relationship between enhanced reward-cue salience and improved successful reversal learning. We speculate that altering sign- and goal-tracking behavior might improve cognitive flexibility in OCD patients, although further research is needed. Our results might explain why WT and reversing SAPAP3^-/-^ (both employing sign tracking and goal tracking) displayed reversal learning, while impaired SAPAP3^-/-^ (refraining from sign tracking) failed the reversal.

### SSRI treatment

Serotonin release in the brain plays a role in behavioral flexibility^32^, and depletion of serotonin leads to impaired reversal-learning performance^33^. Interestingly, in animal models for depression (behaviorally or pharmacologically induced), which show cognitive deficits, both acute and chronic SSRI treatments reversed the cognitive deficits^35,38^. The effects of SSRI treatment on cognitive flexibility in OCD patients have, to our knowledge, not formerly been explored, but a limited number of studies have reported impaired cognitive flexibility in OCD patients with no differences between medicated and unmedicated patients^72–74^. We hypothesized that chronic treatment of SSRI would enhance behavioral flexibility in SAPAP3^-/-^. However, although treatment was effective in reducing excessive grooming and anxiety (as previously reported^44^), it did not alter behavioral flexibility. Taken together, chronic SSRI treatment did not alter behavioral flexibility in SAPAP3^-/-^, a result that may be of clinical relevance for the treatment of OCD.

### Heterogeneity in SAPAP3^-/-^ population

The entire SAPAP3^-/-^ population did not exhibit deficits in behavioral flexibility in our task. Notably, compulsive-like grooming did not correlate with learning or reversal performance, indicating that the severity of compulsive-like grooming per se was not a predictor of behavioral inflexibility. However, we identified half of the SAPAP3^-/-^ population as behaviorally inflexible, demonstrating a striking heterogeneity within the population. Impaired SAPAP3^-/-^ showed normal learning performance, but specifically failed to update previously established cue-reward contingencies. Previous work in OCD reported deficits in behavioral flexibility in patients^10–14^, whereas other studies did not find deficits^15–17^. Benzina and colleagues have identified a subgroup of OCD patients (checking patients) that specifically failed to adjust behavior upon reversal of cue-reward contingencies, and this subgroup was also present in the SAPAP3^-/-^ population^18^. In addition, Manning and colleagues described behavioral deficits in a subgroup of the SAPAP3^-/-^ population, which was independent of compulsive-like grooming levels, similar to our results^47^. Since the SAPAP3^-/-^ in these studies are genetically uniform (between the subgroups of mice that can reverse or are impaired), these results suggest that the differences between the subgroups could be of epigenetic or environmental origin. Lack of a linear relationship between compulsive-like grooming and behavioral flexibility, a clinically relevant finding, suggests that these processes may be driven by independent mechanisms, although we cannot exclude non-linear interactions. Together, these data suggest that a subgroup of SAPAP3^-/-^, similarly to OCD patients, shows diminished behavioral flexibility, which was not associated with compulsive behavior severity.

## Conclusion

In summary, we report that 50 % of the SAPAP3^-/-^ exhibit deficits in behavioral flexibility, independent of compulsivity severity. Impaired SAPAP3^-/-^ refrain from sign-tracking behavior and exclusively rely on goal tracking. Enhancing reward-cue salience restores balanced sign- and goal-tracking behavior, which rescues behavioral flexibility. Thus, our work sheds light on balanced sign and goal tracking that underlies reversal learning and provides a means to enhance behavioral flexibility.

## Acknowledgments

We thank Ralph Hamelink and Dr. Nicole Yee for technical assistance and animal breeding, and Andres de Groot and Mike Vink for technical support. We acknowledge support from the Gravitation program of the Dutch Research Council grant, BRAINSCAPES (024.004.012), and a research grant from FFOR, the Foundation for OCD Research (I.W.).

## Data availability

The data that support the findings in this article and the statistical analyses generated from the data are available on Open Science Framework (https://osf.io/p9k58/).

