## Supplementary information for "Rescuing behavioral flexibility in a mouse model for OCD by enhancing reward-cue salience"

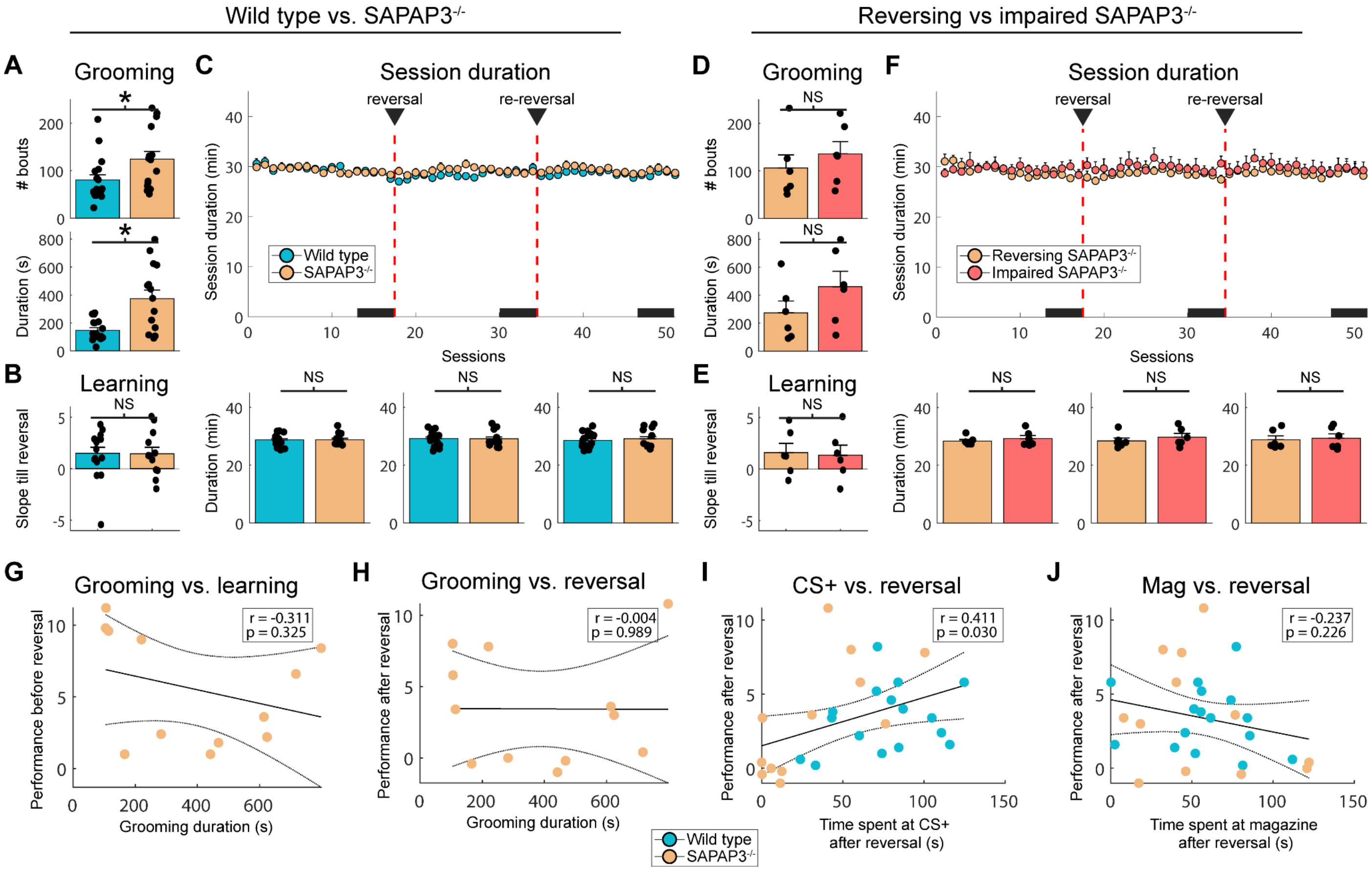


**Supplementary Figure 1.** Supporting data for the regular light-salience experiment. **(A)** Compared to WT (blue, n=17), SAPAP3$^{-/-}$ (orange, n=15) exhibited more grooming bouts (top) and spent more time (bottom) grooming. **(B)** No difference in learning rate between WT and SAPAP3$^{-/-}$ was found. **(C)** Session duration did not differ between WT and SAPAP3$^{-/-}$ (top), as quantified before reversal, after reversal, and after re-reversal (below). **(D)** No difference in number of grooming bouts (top) and grooming duration (bottom) between reversing (orange, n=6) and impaired (red, n=6) SAPAP3$^{-/-}$ was found. **(E)** Learning rate did not differ between reversing SAPAP3$^{-/-}$ and impaired SAPAP3$^{-/-}$. **(F)** Session duration was not different between reversing SAPAP3$^{-/-}$ and impaired SAPAP3$^{-/-}$ (top), neither before reversal, after reversal, or after re-reversal (bottom). **(G)** SAPAP3$^{-/-}$ compulsive-like grooming (n=12) did not correlate with performance before reversal. **(H)** SAPAP3$^{-/-}$ grooming did not correlate with reversal learning. **(I)** Sign tracking of WT and SAPAP3$^{-/-}$ after reversal correlated positively with performance after reversal. **(J)** Goal tracking after reversal did not correlate significantly with performance after reversal.
